# The papain-like protease of coronaviruses cleaves ULK1 to disrupt host autophagy

**DOI:** 10.1101/2020.10.23.353219

**Authors:** Yasir Mohamud, Yuan Chao Xue, Huitao Liu, Chen Seng Ng, Amirhossein Bahreyni, Eric Jan, Honglin Luo

## Abstract

The ongoing pandemic of COVID-19 alongside the outbreaks of SARS in 2003 and MERS in 2012 underscore the significance to understand *betacoronaviruses* as a global health challenge. SARS-CoV-2, the etiological agent for COVID-19, has infected more than 29 million individuals worldwide with nearly ~1 million fatalities. Understanding how SARS-CoV-2 initiates viral pathogenesis is of the utmost importance for development of antiviral drugs. Autophagy modulators have emerged as potential therapeutic candidates against SARS-CoV-2 but recent clinical setbacks underline the urgent need for better understanding the mechanism of viral subversion of autophagy. Using murine hepatitis virus-A59 (MHV-A59) as a model *betacoronavirus*, time-course infections revealed a significant loss in the protein level of ULK1, a canonical autophagy regulating serine-threonine kinase, and the concomitant appearance of a possible cleavage fragment. To investigate whether virus-encoded proteases target this protein, we conducted *in vitro* and cellular cleavage assays and identified ULK1 as a novel bona fide substrate of SARS-CoV-2 papain-like protease (PL^pro^). Mutagenesis studies discovered that ULK1 is cleaved at a conserved PL^pro^ recognition sequence (LGGG) after G499, separating its N-terminal kinase domain from the C-terminal substrate recognition region. Consistent with this, over-expression of SARS-CoV-2 PL^pro^ is sufficient to impair starvation-induced canonical autophagy and disrupt formation of ULK1-ATG13 complex. Finally, we demonstrated a dual role for ULK1 in MHV-A59 replication, serving a pro-viral functions during early replication that is inactivated at late stages of infection. In conclusion, our study identified a new mechanism by which PL^pro^ of *betacoronaviruses* induces viral pathogenesis by targeting cellular autophagic pathway (Word count=250)

**IMPORTANCE:** The recent COVID-19 global pandemic alongside the 2003 SARS and 2012 MERS outbreaks underscore an urgent need to better understand *betacoronaviruses* as pathogens that pose global challenge to human health. Studying the underlying biology of how *betacoronaviruses* subvert innate cellular defense pathways such as autophagy will help to guide future efforts to develop anti-viral therapy. (Word count= 55)

## INTRODUCTION

The recent pandemic of coronavirus disease-2019 (COVID-19) highlights the health crisis worldwide. Although global efforts to restrict travelling and implement social distancing practices have mitigated the spread of Severe Acute Respiratory Syndrome-Coronavirus-2 (SARS-CoV-2), the etiological agent of COVID-19, the virus remains a significant health threat with over 29 million confirmed cases and nearly 1 million fatalities.

Autophagy-modulating drugs such as chloroquine/hydroxychlorine have emerged as potential anti-viral agents against SARS-CoV-2; however, recent placebo-controlled trials showing no clinical benefits and possible safety concerns are spearheading global research efforts for better understanding (1). Autophagy is an evolutionarily conserved process that acts to recycle cellular waste while also responding to invading pathogens. Cellular membranes termed phagophores first enwrap cargo inside double-membraned chambers, so called autophagosomes, after which cargo is degraded upon fusion of autophagosomes with digestive lysosomes. The process is tightly regulated and responsive to various cellular stressors including nutrient-stress, oxidative stress, as well as viral infection (2).

The serine/threonine unc-51-like kinase (ULK1) is a critical upstream regulator of autophagy. The role of ULK1 as a nutrient-responsive orchestrator of autophagy has been well characterized (3). ULK1 is recruited to sites of autophagosome biogenesis where it phosphorylates key autophagy regulatory proteins. Its central role in autophagy has implicated ULK1 in diverse human diseases from cancer and neurodegeneration to inflammatory disorders (4–6). Structurally, ULK1 possesses an N-terminal kinase domain and a C-terminal early autophagy targeting (EAT) domain. The latter facilitates interaction of ULK1 with its various substrates. Autophagy-independent functions of ULK1 have also emerged including the regulation of ER-Golgi trafficking as well as innate immune signaling (7–10). For example, stimulator of interferon genes (STING), a critical adaptor of the DNA sensor cyclic GMP-AMP synthase (cGAS), was previously identified as a substrate of ULK1 (11). Furthermore, the master innate immune kinase TANK binding kinase 1 (TBK1) was reported to be phosphorylated by ULK1 and participate in metabolic signaling (12).

Many positive-sense RNA viruses, including *betacoronaviruses*, have evolved strategies to co-opt autophagy by initiating and utilizing cellular double membrane vesicles as topological surfaces for viral RNA synthesis (13–16); however, the precise mechanisms of viral subversion of autophagy components are poorly defined. The current study uncovers the *betacoronavirus*-encoded papain-like protease (PL^pro^) as a pathogenic factor that disrupts the regulated process of autophagy in part by targeting the autophagy regulatory kinase ULK1.

## RESULTS

### Protein expression of ULK1 is decreased while RNA level is upregulated following mouse coronavirus (M-CoV) infection

To understand the molecular underpinnings of *betacoronavirus* subversion of cellular autophagy, we utilized mouse hepatitis virus-A59 (MHV-A59) as a model and examined its effects on critical components of the canonical autophagy pathway. The serine-threonine kinase ULK1 is an important regulator of cellular autophagy and has recently emerged as an innate immune signaling factor (3, 7–9). The significance of autophagy and innate immune signaling as critical facets of host anti-viral defense, prompted us to investigate the regulation and function of ULK1 during *betacoronavirus* infection. A time-course infection was conducted with MHV-A59 at a multiplicity of infection (MOI) of 10 in the murine fibroblast cell line 17Cl1. Viral replication was verified by immunoprobing for viral non-structural protein 9 (NSP9) precursor. Coincident with active viral polyprotein processing of NSP9 precursor, we observed a significant loss of ULK1 protein starting at 12h post-infection with both anti-ULK1 antibodies tested in this study (**Figure 1A & B**). Interestingly, using the polyclonal anti-ULK1 antibody that recognizes the region of amino acids 351-400, we observed an additional band at ~65 kDa (**Figure 1A**). However, this potential cleavage fragment was undetected with the monoclonal anti-ULK1 antibody that was raised against amino acids 511-750 (**Figure 1B**), indicating a cleavage event may occur within this region. To investigate whether virus-mediated loss of ULK1 is through transcriptional regulation, we performed RT-qPCR. **Figure 1C** showed that mRNA levels of *Ulk1* were elevated following 12h and 24h infection, suggesting that reduced protein expression of ULK1 is not a result of decreased mRNA level. In contrast, elevated gene expression may indicate a compensatory upregulatory mechanism following reduced protein expression.

**Figure 1.**
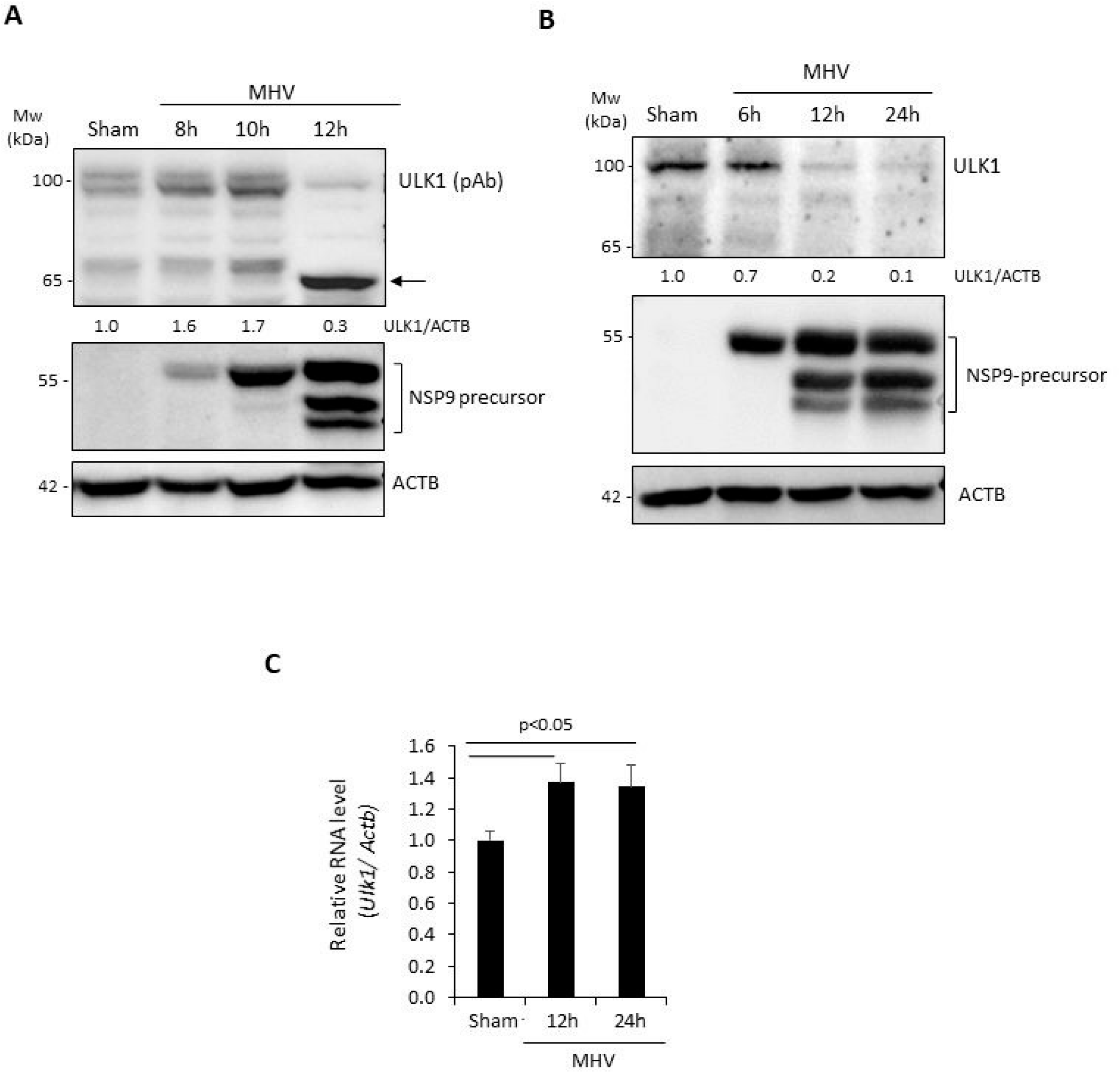
Protein level of ULK1 is decreased following MHV-A59 infection. (A-B) Murine 17Cl1 fibroblasts were infected with MHV-A59 (MOI=10) for various times as indicated or sham-infected with DMEM. MHV replication was assessed *via* western blot analysis of NSP9 precursor processing. Cell lysates were harvested and probed for ULK1 using polyclonal anti-ULK1 (A, which recognizes amino acids 351-400) or monoclonal anti-ULK1 (B, which was raised against amino acids 511-750), NSP9, and ACTB. Densitometric analysis of ULK1 protein level was performed with NIH ImageJ, normalized to ACTB and presented underneath as fold changes where the first lane was arbitrarily set to 1. Arrow denotes the cleavage fragment detected at ~65 kDa. (C) Murine 17Cl1 cells were sham-infected or infected with MHV-A59 (MOI=10) for 12h or 24h. Gene level of *Ulk1* was measured by real-time quantitative RT-PCR and normalized to *Actb* (mean ± SD, n=3).

### PL^pro^ cleaves ULK1

The discovery of a potential ULK1 cleavage product in MHV-infected lysates prompted us to investigate whether *betacoronaviral* proteases are responsible for ULK1 cleavage. *Betacoronaviruses* encode two proteases, a papain-like cysteine protease (PL^pro^) and a 3-chymotrypsin-like cysteine protease (3CL^pro^, also known as the Main protease, M^pro^) that process viral polyproteins into individual functional proteins (17). We generated a construct expressing PL^pro^ of SARS-CoV-2, which has 63% similarity with PL^pro^ domain 2 of MHV-A59 (**Figure 2A**). To determine whether PL^pro^ targets ULK1, HEK293T cells were transiently transfected with either a control vector or a plasmid expressing PL^pro^ for 24h. Expression of PL^pro^ alone was sufficient to recapitulate the reduction in full-length ULK1 observed following MHV-A59 infection using the monoclonal anti-ULK1 antibody (**Figure 2B**). PL^pro^-mediated cleavage of endogenous ULK1 was then confirmed using N-terminal 3×Flag-ULK1 construct (**Figure 2C**). HEK293T cells were co-transfected with PL^pro^ together with 3×Flag-ULK1. Lysates probed with an anti-Flag antibody revealed a significant reduction of ULK1 with the concomitant detection of a lower-molecular-weight fragment of ~70kDa in size. To exclude the potential involvement of 3CL^pro^, we performed in *vitro* cleavage assay. Time course treatment revealed that ULK1 was not targeted by the SARS-CoV-2 3CL^pro^, whereas coxsackievirus 3C^pro^, the positive control, demonstrated an efficient cleavage of ULK1 **(Figure 2C)**. Together, our data suggest that PL^pro^, but not 3CL^pro^, targets ULK1 for cleavage.

**Figure 2.**
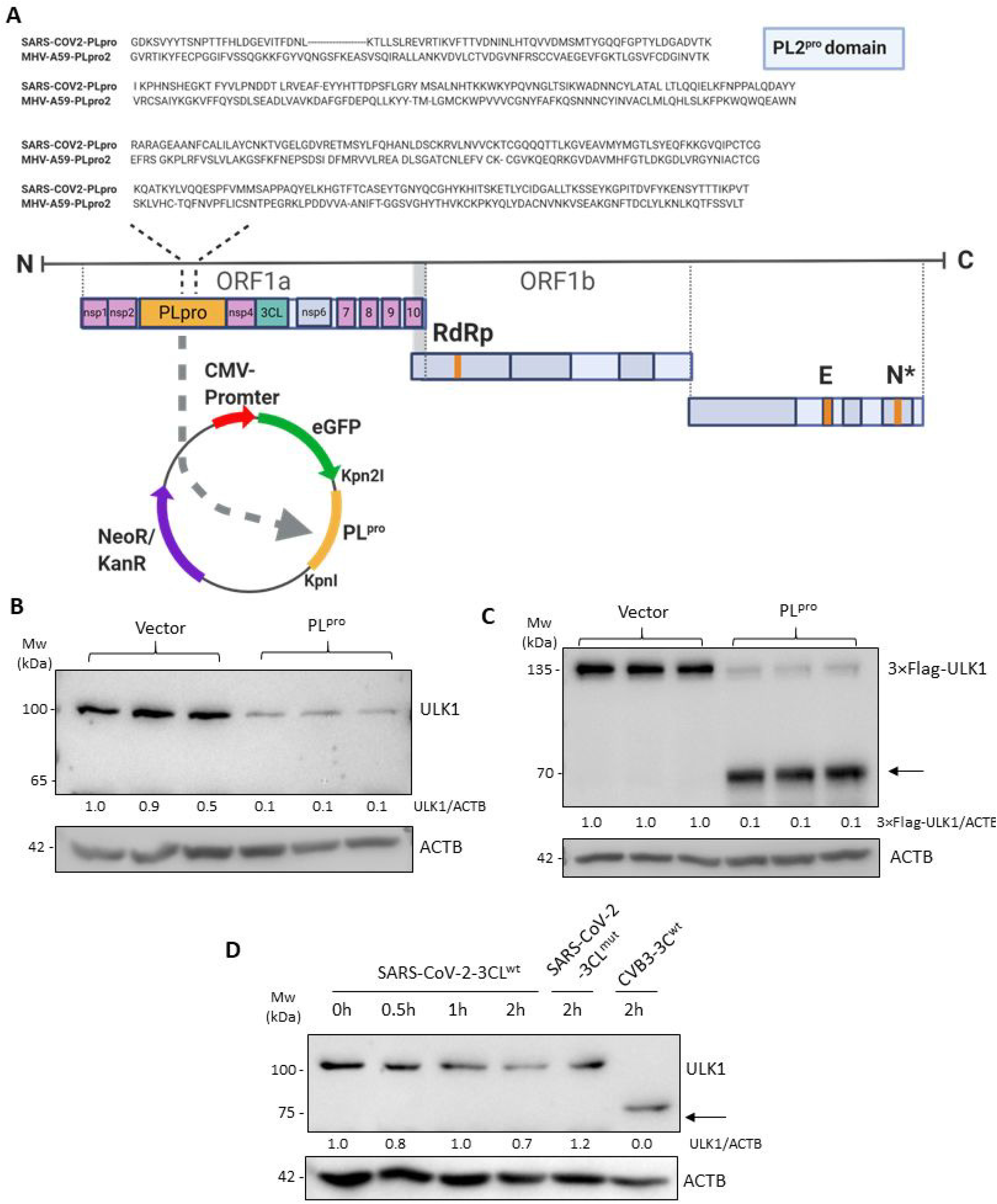
PL^pro^ of SARS-CoV-2 cleaves ULK1. (A) Sequence alignment of the PL protease domain of SARS-CoV-2 and PL protease domain 2 of MHV-A59 is shown. The SARS-CoV2-PL^pro^ was cloned into EGFP-C1 vector. (B) HEK293T cells were transfected with either control or PL^pro^-expressing plasmid for 24h. Lysates were probed by western blotting for endogenous ULK1 using the monoclonal anti-ULK1 antibody and normalized to ACTB as in Figure 1A. (C) HEK293T cells were co-transfected with 3×Flag-ULK1 and either control vector or PL^pro^-expressing plasmid for 24h. Lysates were probed with anti-Flag antibody to detect exogenous ULK1. Arrow denotes cleavage fragment observed at ~70 kDa. Densitometry is provided as in Figure 1A. (D) Purified SARS-CoV-2 3CL^wt^ (4 μg) or catalytically-inactive C145A 3CL^mut^ (4 μg) of SARS-CoV-2 and CVB3 3C^wt^ (0.1 μg) alongside HeLa lysates (30 μg) were used to perform *in vitro* cleavage assay. Western blotting was performed to probe for endogenous ULK1 and normalized to ACTB as in Figure 1A. Arrows denote cleavage fragments.

### PL^pro^ cleaves ULK1 after G499

To identify the precise location within ULK1 that is targeted by PL^pro^, we closely investigated the protein sequence of ULK1 and identified two potential cleavage sites in the central region of ULK1 with a previously reported consensus cleavage motif (**Figure 3A)** (18). Using the Flag-tagged ULK1 construct, we mutated the potential cleavage sites and co-transfected wild-type (WT) or mutant ULK1 with either control or PL^pro^ construct into HEK293T cells. Expression of PL^pro^ resulted in the loss of the 3×Flag-ULK1 and the appearance of a faster migrating band at ~70 kDa in cells expressing WT-ULK1 and G531 mutant ULK1, but not in cells expressing G499 mutant ULK1(**Figure 3B)**, indicating that PL^pro^ cleaves ULK1 after G499. The observed N-terminal fragment matches the molecular weight of the predicted cleavage fragment. The resulting cleavage of ULK1 by PL^pro^ separates its N-terminal kinase domain from a C-terminal substrate binding domain (**Figure 3C**).

**Figure 3.**
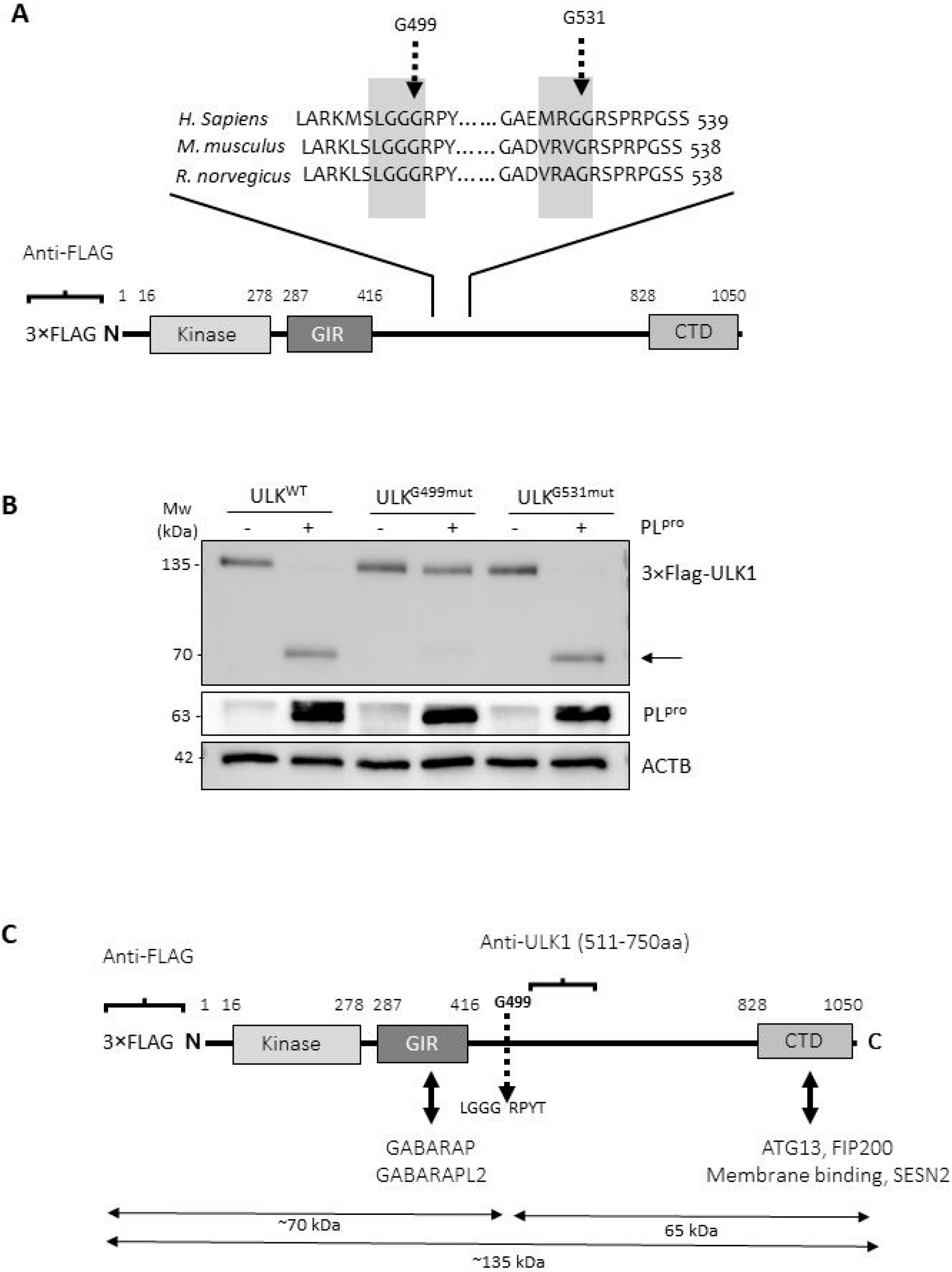
PL^pro^ cleaves ULK1 after G499. (A) PL^pro^ consensus cleavage motif in ULK1 from H. sapiens, M. musculus, and R. norvegicus is presented. Highlighted gray boxes denote potential consensus cleavage sites. Dashed arrows denote sites of potential cleavages. (B) HEK293T cells were co-transfected with PL^pro^ and either 3×Flag-ULK1^WT^, 3×Flag-ULK1-G499^mut^ or 3×Flag-ULK1-G531^mut^. Western blotting was conducted with anti-Flag and anti-GFP antibody for detection of ULK1 and PL^pro^, respectively. (C) Schematic diagram of ULK1 protein sequence with corresponding functional domains, antibody recognition epitopes and identified PL^pro^ cleavage site.

### PL^pro^-mediated cleavage of ULK1 disrupts autophagy

To understand the functional consequence of PL^pro^-mediated cleavage of ULK1, we first examined whether the ability of ULK1 to interact with its binding partners is disrupted. ATG13 is a bridging molecule that directly interacts with both ULK1 and the scaffold protein FIP200 in the autophagy-initiating ULK1 complex (3). Expression of PL^pro^ resulted in reduced coimmunoprecipitation of ULK1 with its binding partner ATG13 (**Figure 4A)**. Given that the ULK1 complex is required for starvation-induced canonical autophagy, we inquired the functional significance of its disruption. The rate of autophagic degradation, designated as ‘autophagy flux’, can be measured by comparing levels of LC3-II, a substrate of autophagy, in the presence or absence of a lysosomal fusion inhibitor (19). 17Cl1 cells were sham-or MHV-A59-infected for 6h, 12h, and 24h either in the presence of DMSO or a lysosomal fusion inhibitor, bafilomycin A1 (BAF). Consistent with impaired autophagy, MHV-infected cells had a reduced accumulation of LC3-II in the presence of BAF (**Figure 4B**). We next tested whether PL^pro^ expression can recapitulate the impaired autophagy observed following MHV-A59 infection. HEK293T cells were transfected with either a control vector or a plasmid expressing PL^pro^ for 24h and then subjected to starvation in HBSS medium for 2h in the presence or absence of BAF. Control cells responded efficiently to starvation by increasing autophagy flux; however, cells expressing PL^pro^ did not initiate autophagy (**Figure 4C**), suggesting that PL^pro^ expression disrupts cellular autophagy.

**Figure 4.**
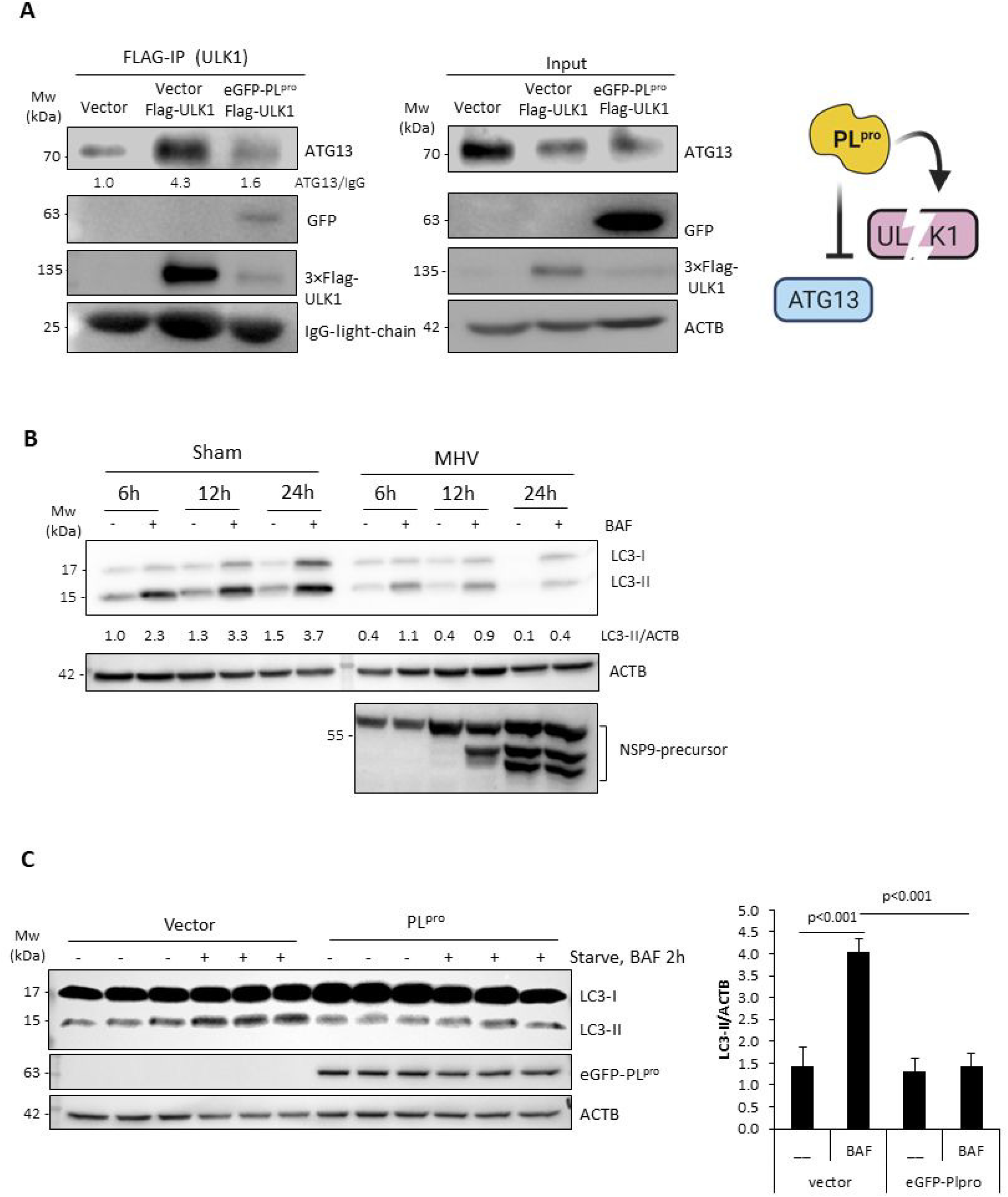
PL^pro^-mediated cleavage of ULK1 disrupts autophagy. (A) HEK293T cells were co-transfected with 3×Flag-ULK1 together with control vector or PL^pro^ for 24h. Immunoprecipitation of exogenous 3×Flag-ULK1 was performed using anti-Flag M2 agarose beads. IgG light chain and ACTB served as loading controls for immunoprecipitation and inputs, respectively. Schematic depiction of disrupted formation of ULK1-ATG13 complex (right). (B) Disruption of autophagy flux in MHV-A59-infected cells. Autophagy flux was assayed in the presence or absence of BAF (125 nM) following sham or viral infection. Cell lysates were analyzed by western blotting for LC3 and normalized to ACTB. NSP9 was used as marker of active viral replication. (C) HEK293T cells expressing either empty vector or PL^pro^ construct were starved for 2h in the presence or absence of lysosomal inhibitor BAF (125 nM) for 2h. Lysates were probed for LC3 and normalized to ACTB. Quantification of LC3-II was conducted as in Figure 1A and presented as (mean ± SD, n=3) in the right panel.

### ULK1 plays a dual role in MCoV replication

The role of ULK1 in *betacoronavirus* replication is currently unclear. To clarify this, we utilized chemical and genetic approaches to target ULK1 prior to infection with MHV-A59. The chemical inhibitor, MRT68921, potently targets ULK1/2 kinase activity (20). Functional validation of the inhibitor showed the complete blockage of starvation-induced autophagy (ie. reduced LC3-II accumulation in starvation-induced, BAF-treated cells **(Figure 5A)**. To investigate the consequence of ULK1 inhibition, 17Cl1 cells were inoculated with MHV-A59 for 1h, followed by replenishment with medium containing either DMSO or MRT68921 (5 μM). After 12 or 24h, the *N-gene* levels of MHV-A59 were measured by RT-qPCR. Compared to control treatment, MRT68921 significantly attenuated viral RNA replication at 24h post infection (**Figure 5B**). Consistent with a potential pro-viral role for ULK1, RNA levels and viral titers of MHV-A59 were significantly reduced following gene-silencing of *Ulk1* (**Figure 5C**). We also assessed the effects of expression of WT- or non-cleavable mutant-ULK1 on viral replication. We showed that cells expressing WT-ULK1 displayed significantly enhanced viral titers (**Figure 5D**), supporting a pro-viral function for ULK1. However, a non-cleavable ULK1 mutant demonstrated diminished viral titers (**Figure 5D**). Collectively, these data support a requirement of ULK1 for replication prior to its cleavage.

**Figure 5.**
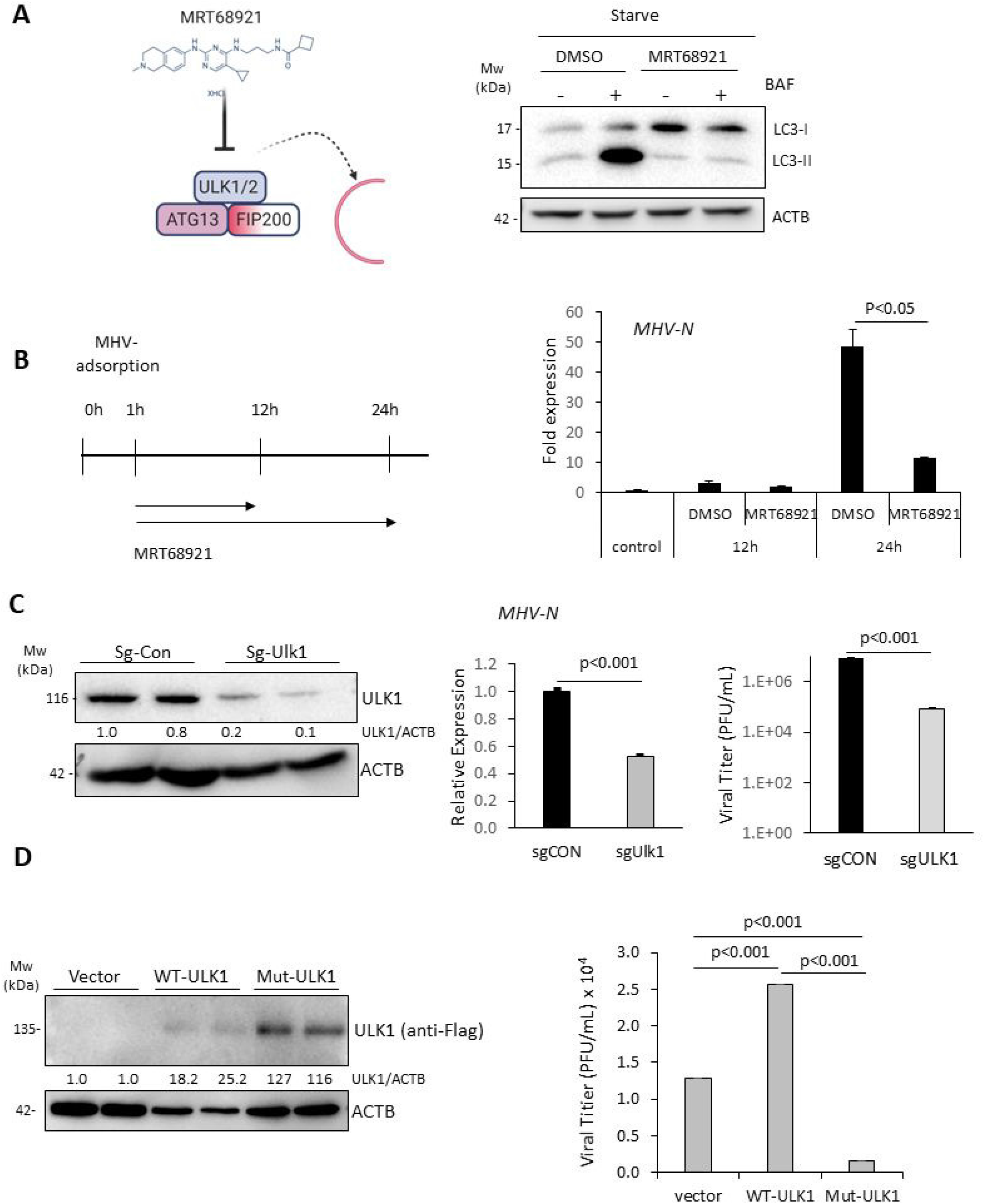
ULK1 plays a dual role in MHV-A59 replication. (A) Schematic diagram of the chemical structure and mechanism of action of MRT68921 (left). HEK293T cells were starved for 2h in the presence or absence of MRT68921 (5 μM) and BAF (125 nM) as indicated. Western blotting was carried out for detection of LC3 and ACTB. (B) Schematic illustration of MRT68921 treatment of 17Cl1 cells following MHV-A59 infection (left). 17Cl1 cells were infected with MHV-A59 (MOI=10) for indicated time-points in the presence of DMSO (vehicle control) or MRT68921 (5 μM). Relative MHV-*N* RNA level was determined by RT-qPCR and normalized to *Actb* (mean ± SD, n=3). (C) Murine 17Cl1 cells were transfected Cas9 and sg-CON or sg-ULK1 for 48h. Cells were then infected with MHV-A59 (MOI=10) for an additional 24h. ULK1 knockout efficiency (left), viral RNA levels in the cells (middle) and viral titers in the supernatant (right) were measured by western blotting, RT-qPCR, and TCID50 assay, respectively. (D) 17Cl1 cells were transfected with vector, 3× Flag-ULK1^WT^, or 3× Flag-ULK1^MUT^ for 24h. Cells were then infected with MHV-A59 (MOI=10) for 24h. Cell lysates were analyzed by western analysis for exogenous ULK1 with anti-Flag antibody. Viral titers in culture medium were determined by TCID50 assay (mean ± SD, n=3).

## DISCUSSIONS

Diverse viruses have evolved strategies to circumvent the inherently anti-viral defense capacity of autophagy (21). In particular, RNA viruses that replicate in the cytoplasm can utilize autophagic membranes as topological surfaces for viral replication (15, 22, 23). However, viruses must evade selective targeting and degradation via autophagy, termed virophagy, a process mediated by autophagy receptors that recognize and sequester viral components inside autophagosomes (24). The autophagy receptor SQSMT1 was previously shown to mediate virophagy of the positive-strand RNA virus Sindbus (25). To counteract this, enterovirus such as coxsackievirus B3 and enterovirus-D68 have evolved strategies to subvert the host virophagy efforts by engaging virus-encoded proteases to cleave SQSTM1 (26, 27). Similarly, enteroviral proteases impede the degradation capacity of autophagy by disrupting autophagosome-lysosome fusion through the selective cleavage of autophagic fusion SNARES (19, 28). Recent studies are beginning to unravel the subversion strategies utilized by other RNA viruses. Global outbreaks of *betacoronaviruses* in 2003 (SARS-CoV-1), 2012 (MERS), and 2019 (SARS-CoV-2) have spearheaded efforts to better understand their relationship with autophagy. Despite great efforts, there is a limited understanding of whether *betacoronavirus*-encoded proteases target host substrates.

The murine *betacoronavirus* MHV-A59 was reported to induce double membrane vesicles reminiscent of autophagosomes in the absence of intact autophagy (29), while infection of porcine *betacoronavirus,* PHEV, demonstrated significant reduction of ULK1 (30). Findings from the current study suggest that *betacoronaviruses* may actively target canonical autophagy factors. We provide evidence that SARS-CoV-2-endoced PL^pro^ can cleave ULK1 to disrupt formation of the autophagy-initiating ULK1-complex and functionally impair starvation-induced cellular autophagy/degradative capacity. SARS-CoV-1 PL^pro^ was previously reported to have de-ubiquitinase activity by recognizing the consensus LXGG motif similar to other host deubiquitinating enzymes (31). Of note, we identified the SARS-CoV-2 PL^pro^ cleavage site of ULK1 after glycine (G) 499, follows precisely the consensus sequence recognized by the de-ubiquitinase. The interferon regulatory factor 3 (IRF3) was also recently reported as a substrate of PL^pro^ following an LGGG consensus motif similar to ULK1 (32). The targeting of ULK1 and possibly other host proteins harboring an LXGG motif therefore underscores potentially a novel mechanism through which *betacoronaviruses* subvert cellular autophagy. In addition to disrupting ULK1-mediated autophagy, the deubiquitinase activity of PL^pro^ may favor viral pathogenesis by disrupting selective autophagy, a process that relies on autophagy receptors recognizing ubiquitin-modified pathogens or cellular cargo (33).

The role of ULK1 in *betacoronaviral* replication is poorly defined. We found ULK1 protein levels are relatively intact during the initial 6h of MHV infection. Furthermore, loss-of-function studies prior to infection either through chemical inhibition of ULK1/2 kinase activity or genetic silencing of ULK1 revealed a significant reduction in MHV viral replication. In contrast, ULK1 protein levels are significantly decreased during late infection, coinciding with the emergence of lower-molecular-weight cleavage fragment. Additionally, the expression of WT-ULK1 that can undergo proteolytic processing by PL^pro^ enhances viral titers unlike the expression of a non-cleavable ULK1. Collectively, these data suggest that ULK1 may serve a hitherto uncharacterized pro-viral role during early replication. The cleavage of ULK1 during late infection likely suggests that (1) ULK1 function may be dispensable at late stages; (2) ULK1 may serve late-stage anti-viral function that needs to be inactivated; and/or (3) ULK1 cleavage fragments may serve novel pro-viral function. The attenuation of viral titers in cells expressing non-cleavable ULK1 suggests that persistent ULK1 activity throughout the course of infection may be undesirable for virus. Indeed, ULK1 has previously been reported to regulate anti-viral innate immune signaling (11, 12).

In summary, we uncover a novel function for the PL^pro^ of SARS-CoV-2 in cleaving the autophagy-regulating kinase ULK1. These insights clarify the mechanism behind *betacoronaviral* pathogenesis and subversion of cellular autophagy.

## MATERIALS AND METHODS

### Cell culture, viral infection, and chemicals

Murine 17Cl1 fibroblast cells were cultured in Dulbecco’s Modified Eagle’s Medium (DMEM) supplemented with 10% fetal bovine serum (FBS) and a penicillin/streptomycin cocktail (100 μg/mL). MHV-A59 and 17Cl1 cells were provided by Dr. Nerea Irigonen (University of Cambridge).

For MHV-A59 infection, cells were either sham-infected with DMEM or inoculated with MHV-A59 at a multiplicity of infection (MOI) of 10. Cells were starved by culturing in Hank’s Balanced Salt Solution (HBSS) medium (Thermofisher Scientific, 14025076) for 2h. The V-ATPase inhibitor bafilomycin A1 (BAF, Sigma-Aldrich, B1793) was used at a concentration of 125 nM. The ULK1/2 kinase inhibitor MRT68921 HCl (Selleck, S7949) was used at a dose of 5 μM.

### Plasmids

The 3×Flag-ULK1 plasmid was generated using a multiple cloning site modified CMV10 vector backbone with the corresponding cut sites for ULK1 (EcorI/BamHI). Cleavage resistant mutants, LGGG-499-QQQG and MRGG-531-MRDD, were generated using gBLOCKS DNA synthesis (IDT) and fragments were cloned through restrict digestion of ULK1 with BstEII/FseI and FseI/AflII, respectively. The cloning strategy for eGFP-PL^pro^ of SARS-CoV-2 is outlined in **Figure 2A**. Briefly, PL^pro^ sequence of SARS-CoV-2 was obtained from NCBI [Wuhan Hu-1 Isolate reference sequence: NC_045512.2] and generated using gBLOCKS DNA synthesis (IDT) with nucleotides corresponding to residues 713-1063 of SARS-CoV-2 PL^pro^ sequence and cloned into N1-pEGFP vector at Kpn2 and KpnI sites. The single guide RNA (sgRNA) sequence targeting murine *Ulk1* is AATCTTGACTCGGATGTTGC. For transfection, cells were transiently transfected with plasmid cDNAs or sgRNAs using Lipofectamine 2000 (Invitrogen, 11668-019) following the manufacturer’s instructions.

### Purification of SARS-CoV-2 3CL^pro^

pET-41c plasmids encoding either wild-type (WT) or mutant (C145A) SARS-CoV-2 3CL^pro^ were transformed into C41 (DE3) *E. coli* and then plated onto kanamycin (50 μg/mL) agar plates. A starter culture from a single colony was grown overnight and then diluted 100-fold in Terrific Broth [Sigma, T9179]. Expression was induced with 1mM isopropyl ß-D-1-thiogalactopyranoside (IPTG) after cultures reached an OD600 of 0.6-0.8 and proceeded at 25°C for 5 additional hours. Protein was purified using Ni-NTA Fast Start (Qiagen, #30600) according to the manufacturer’s instructions.

### *In vitro* cleavage assay

*In vitro* cleavage assay was performed as previously described (27). Briefly, HeLa lysates (20 μg) were incubated with WT or catalytically inactive (C145A) SARS-CoV-2 3CL^pro^ (4 μg) in cleavage assay buffer (20 mM HEPES pH 7.4, 150 mM KOAc, 1 mM DTT) for the indicated times at 37°C. Reactions were terminated with 6×sample buffer and subjected to western blot analysis.

### Western blot analysis

Cells were lysed in buffer (10 mM HEPES pH 7.4, 50 mM NaPyrophosphate, 50 mM NaF, 50 mM NaCl, 5 mM EDTA, 5 mM EGTA, 100 μM Na_3_VO_4_, 0.1% Triton X-100) and western blotting was conducted using the following primary antibodies: LC3 (Novus Biologicals, NB100-2220), actin-*beta* (ACTB, Sigma-Aldrich, A5316), monoclonal ULK1 (Santa Cruz, sc-390904), polyclonal ULK1 (Abcam, ab167139), nonstructural protein 9 (NSP9, Sigma, SAB3701435), FLAG (Sigma, F1804), and GFP (Life Technologies, A-6455).

### *Ex vivo* cleavage assay

*Ex vivo* or cellular cleavage assay was performed as previously described (27). Briefly, HeLa cell were transfected with 3×Flag-ULK1 (WT or mutant) and either control vector or eGFP-PL^pro^ for 16h. Lysates were harvested and subjected to western blot analysis with anti-Flag-antibody to detect cleavage fragments.

### Immunoprecipitation

Immunoprecipitation (IP) of Flag-tagged ULK1 construct was performed using EZview^TM^ Red ANTI-FLAG ^®^ M2 Affinity Gel (Sigma-Aldrich, F2426) according to the manufacturer’s instructions. In brief, HeLa cell lysates were incubated with anti-Flag M2 agarose beads at 4°C overnight. After three washes, the bound proteins were eluted with 2×SDS sample buffer and then subjected to western blot analysis.

### Viral titer measurement

Samples were serially diluted and overlaid on 60-well Terasaki plates of HeLa cells. After 48h incubation, 50% tissue culture infective dose titer (TCID50) was calculated by the statistical method of Reed and Muench (34). Titers were expressed as plaque forming unit (PFU)/mL with 1 infectious unit equal to 0.7 TCID50 as described previously (35).

### Real-time quantitative RT-PCR

Total RNA was extracted using the RNeasy Mini kit (Qiagen, 74104). To determine gene expression levels, quantitative PCR targeting *Ulk1* (forward: GCAGCAAAGACTCCTGTG ACAC; reverse primer: CCACTACACAGCAGGCTATCAG), and *MHV N* gene (forward primer: CAAAGAAAAGGGCGTAGACAGG; reverse primer: CGCCATCATCAAGGATCTGAG) was performed in a 10-μl reaction containing 1 μg of RNA using the TaqMan™ RNA-to-CT™ 1-Step Kit (Life Technologies, 4392653) and normalized to *Actb* mRNA according to the manufacturer’s instructions. The PCR reaction was performed on a ViiA 7 Real-Time PCR System (Applied Biosystems). Samples were run in triplicate and analyzed using comparative CT (2-ΔΔCT) method with control samples and presented as relative quantitation (RQ).

### Statistical analysis

Results are presented as mean ± standard deviation (SD). Statistical analysis was performed with unpaired Student’s *t-*test or analysis of variance (ANOVA). A p-value <0.05 was considered to be statistically significant. All results presented are representative of at least 3 independent experiments.

## ACKNOWLEDGEMENTS

This work was supported by the Natural Sciences and Engineering Research Council (RGPIN-2016-03811), Canadian Institutes of Health Research (PJT-159546 and PJT-173318), and the Heart & Stroke Foundation of Canada (G-18-0022051) to HL. YM is the recipient of a Doctoral Fellowship from ALS Canada-Brain Canada. YM, YCX, and AB are recipients of a four-year PhD Fellowship from the University of British Columbia. YCX is a recipient of the CIHR Doctoral Fellowship. CSN and HTL are supported by the MITACS Accelerate program.

